# Cargo adaptor identity controls the mechanism and kinetics of dynein activation

**DOI:** 10.1101/2024.10.09.617440

**Authors:** John P. Gillies, Saffron R. Little, William O. Hancock, Morgan E. DeSantis

## Abstract

Cytoplasmic dynein-1 (dynein), the primary retrograde motor in most eukaryotes, supports the movement of hundreds of distinct cargos, each with specific trafficking requirements. To achieve this functional diversity, dynein must bind to the multi-subunit complex dynactin and one of a family of cargo adaptors to be converted into an active, processive motor complex. Very little is known about the dynamic processes that promote the formation of this complex. To delineate the kinetic steps that lead to dynein activation, we developed a single-molecule fluorescence assay to visualize the real-time formation of dynein-dynactin-adaptor complexes in vitro. We found that dynactin and adaptors bind dynein independently rather than cooperatively. We also found that different dynein adaptors promote dynein-dynactin-adaptor assembly with dramatically different kinetics, which results in complex formation occurring via different assembly pathways. Despite differences in association rates or mechanism of assembly, all adaptors tested can generate a population of tripartite complexes that are very stable. Our work provides a model for how modulating the kinetics of dynein-dynactin-adaptor binding can be harnessed to promote differential dynein activation and reveals a new facet of the functional diversity of the dynein motor.

## Introduction

Cytoplasmic dynein-1 (dynein) is a 1.5 MDa motor protein complex that uses energy derived from ATP hydrolysis to move processively toward the minus-end of the microtubule.^1^ A single dynein motor complex contains two copies each of six different subunits (Figure 1A).^2^ In all eukaryotes, except land plants, dynein is responsible for most long-distance retrograde cargo trafficking.^3^ Dynein also has essential roles in building and aligning the mitotic spindle and promoting the metaphase-anaphase transition by silencing the Spindle Assembly Checkpoint.^1^ To support these diverse functions, dynein interacts with a network of regulatory proteins. In fact, dynein is unable to bind cargo or move processively on the microtubule unless it is in complex with the multi-subunit complex, dynactin, and one of a family of cargo adaptors.^4^ The complex of dynein-dynactin-adaptor, which we will refer to as the *activated transport complex*, is the functional unit that is responsible for dynein-driven cargo motility (Figure 1A).^5,6^ There are nearly 20 identified adaptors in humans, many of which localize to distinct cargos.^4^ This means that adaptors are responsible, in part, for dynein’s ability to traffic cargo with specificity. How dynein discriminates between adaptors to bind to specific cargos remains an outstanding question.

**Figure 1:**
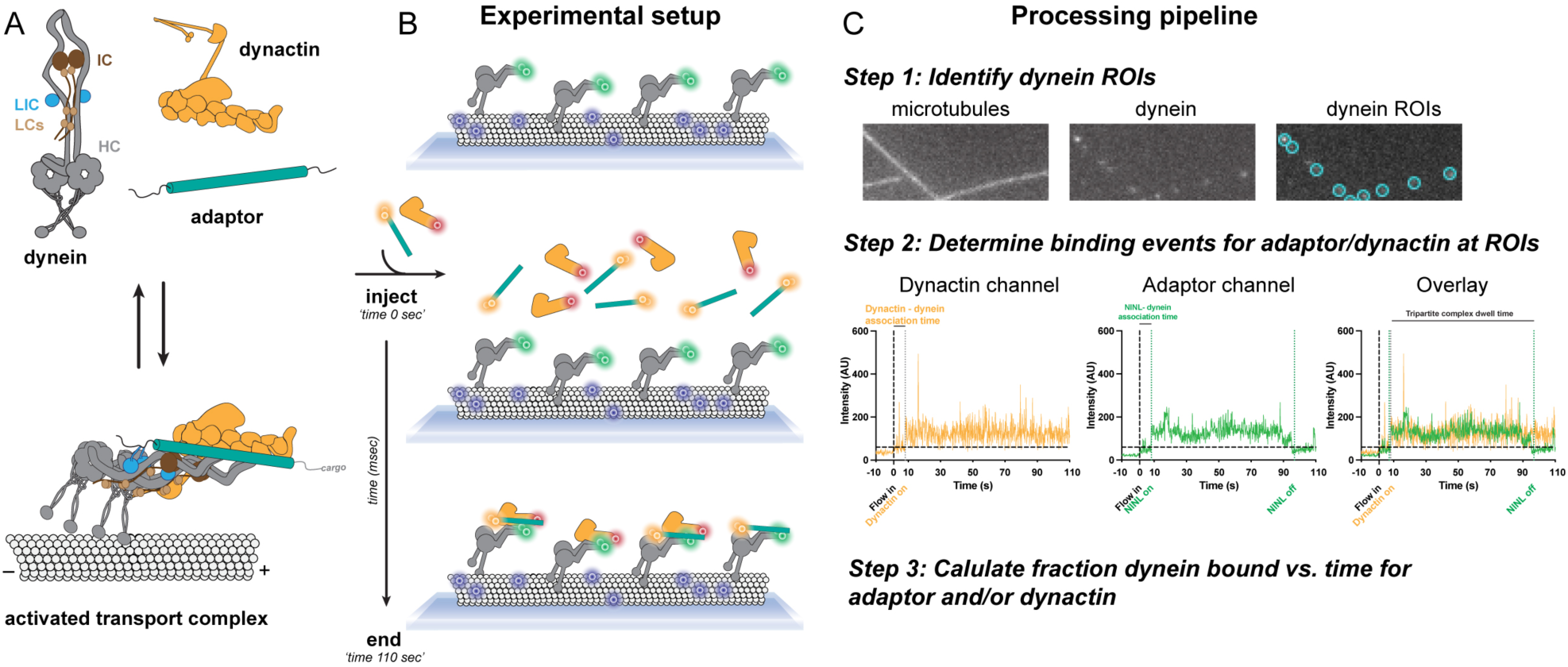
Single molecule association assay development to study dynein binding kinetics. **A.** Diagram of dynein activation. Dynein (heavy chain (HC): gray; intermediate chain (IC): dark brown; light intermediate chain (LIC): blue; light chains (LCs): light brown) must bind to dynactin (yellow) and adaptor (teal) to form an activated transport complex capable of motility on a microtubule. **B.** Experimental setup. Dynein is tightly associated with microtubules, after which dynactin and/or adaptor are flowed in and the intensity of fluorescence is monitored at dynein puncta. **C.** Processing pipeline. ROIs were generated at individual dynein spots. The fluorescence intensity of adaptor and/or dynactin was measured at dynein ROIs. Some kinetic parameters that can be extracted from intensity values (association time and dwell time) are indicated in the figure. These signals were compared to a baseline value to determine when dynactin and adaptor were bound to dynein.

Multiple structural and biochemical studies have revealed how dynactin and an adaptor convert dynein from an autoinhibited conformation, called Phi, to an active and processive motor (Figure 1A).^5–11^ In the Phi conformation, the motor domains are held in a crossed conformation that results in a low affinity, non-productive interaction between dynein and the microtubule track.^10^ In contrast, in the activated transport complex, dynein’s motors are in a parallel conformation, which places both microtubule binding domains in the same orientation and allows the motor domain to productively hydrolyze ATP.^7,8,11^ These conformational changes are driven by extensive interactions that occur between multiple dynein subunits, dynactin subunits, and the adaptor (Figure 1A).

Despite knowing its final structure, we know almost nothing about the dynamic processes that govern how the activated transport complex forms. Elucidating the mechanism of how dynein binds to dynactin and adaptor is essential as dynein activation is the direct outcome of the formation of this complex. To address this gap and uncover the mechanism of how dynein-dynactin-adaptor binding occurs, we developed a single-molecule based assay to monitor complex formation in real time using Total Internal Reflection Fluorescence (TIRF) microscopy. We found that different adaptors assemble into the activated transport complex with different kinetics and via a different mechanism, but that once formed, all adaptors tested can form very stable, long-lived activated transport complexes. We also found that the order of complex assembly can influence the resultant complex stability. These findings suggest that differences in dynein-dynactin-adaptor binding kinetics or assembly mechanisms may contribute to how dynein achieves adaptor binding specificity in cells and helps to explain the diversity of dynein function.

## Results

### Single-molecule association assay development

Most of what we know about dynein’s association with dynactin and adaptors has been obtained via motility experiments that are performed without consideration of the effect of binding kinetics. In these types of studies, low concentrations of dynein, dynactin, and adaptors are mixed, allowed to incubate, and the motility of the resultant complexes is visualized in movies obtained via TIRF microscopy. While this approach is powerful because it allows for the quantification of how the assembled activated transport complexes move on the microtubule track, these motility assays cannot return information about how the activated transport complex forms. Further, while these assays indirectly probe the result of complex formation (motility), they do not directly report on dynein’s affinity for dynactin or adaptor or relay information about the stability of the formed complex. Additionally, traditional motility experiments cannot address whether dynactin or the adaptor must bind first to form the activated transport complex, or whether the identity of different adaptors bias activated complexes to form through different mechanisms of assembly.

The aim of this study was to obtain information about the dynamic processes that underly dynein activation. Specifically, we set out to address two key questions: 1*) Does the activated transport complex assemble via an ordered or random mechanism? 2) Do different adaptors form activated transport complexes with different kinetics or order of assembly?* To answer these questions, we developed a single-molecule assay to visualize dynein-dynactin-adaptor association in real time via TIRF microscopy (Figure 1B and C). First, Alexa-488-labeled dynein was incubated with preassembled Alexa-405-labeled microtubules in a flow chamber (Figure 1B). We included the nucleotide hydrolase apyrase in the binding buffer to ensure that ATP carried over from the purification was depleted, and dynein remained in the apo conformation, which has a high affinity for microtubules. Next, using a custom-built apparatus that allowed us to simultaneously image and inject sample, we flowed in fluorescently labeled dynactin and adaptor and acquired movies of dynactin and adaptor interacting dynamically with dynein at a frame rate of 150 msec/frame (Figure 1B).

To quantify binding kinetics, we monitored dynactin and adaptor intensity over time at dynein puncta, which allowed us to semi-automate the analysis. To accomplish this, we first identified regions of interest (ROIs) as dynein puncta colocalized with microtubules that were present in the first and last frame (Figure 1C- Step 1). We then simultaneously monitored the intensity of the dynactin and/or adaptor fluorophores at the dynein ROIs (Figure 1C- Step 2). Potential binding events were defined as positive deviations in intensity from baseline (as determined after rolling ball background subtraction of each frame), while unbinding events were defined by an elevated intensity returning to baseline (Figure 1C- Step 2). Previous studies have shown that the activated transport complex often contains two dynein dimers and can also contain two adaptor dimers.^7,8^ An experimental limitation of this assay is that it is monitors the association of one dynein dimer with a dynactin complex and one adaptor dimer. For an extended discussion of this caveat, refer to the Methods.

The TMR and JF646 dyes used to label dynactin and adaptor inherently blink and photobleach over time (Figure S1A and B). To increase confidence that transitions between bound and unbound events reflected changes in protein-protein interactions rather than dye blinking, we enforced an 8-frame persistence cutoff, which means that all events had to last longer than 1.2 seconds to be considered a bona fide instance of binding or unbinding (Figure S1C). To reduce the effect of photobleaching on our experimental measurements, we limited the duration of our movies to 120 seconds, a time frame in which greater than 60% of both dyes remained unbleached (Figure S1A). To rule out the possibility that the specific dyes used on dynactin or adaptors affected the kinetics, most experiments in this study were collected with biological replicates where the identity of the dyes on dynactin and adaptor were swapped. Additionally, we found that the time it took dynactin or adaptor to bind dynein in our studies was independent of the identity of the dye. (Figure S1D).

This assay can yield many kinetic parameters. Association rate constants can be extracted from measurements of the fraction of dynein ROIs colocalized with dynactin and/or adaptor over time under different experimental conditions (Figure 1C). By monitoring both dynactin and adaptor channels simultaneously, we can also extract which component associated with dynein first at individual dynein ROIs (Figure 1C). Off-rates can be calculated directly by determining the number of unbinding events divided by total time dynein molecules are bound to dynactin and/or adaptor. Finally, complex stability can also be assessed by measuring the dwell time of complexes formed at each dynein ROI (Figure 1C).

### NINL and dynactin bind dynein independently

We next set out to establish that the single molecule association assay we developed could report on dynein’s association with dynactin and the adaptor NINL, which is proposed to link dynein to MICAL3-RAB8A vesicles and melanosomes.^12,13^ When not bound to cargo, many adaptors assume a folded, autoinhibited conformation in which their C-terminal end folds back to block dynein and dynactin association with the adaptors’ N-terminal coiled coil.^11,14,15^ To circumvent potential adaptor autoinhibition and analyze dynein motility in the absence of cargo, many published motility studies use adaptors with C-terminal truncations. For this study, we used a well-characterized truncation of NINL that robustly activates dynein motility and increases the yield and ease of the NINL protein purification.^16,17^

First, we measured the association kinetics of dynein-NINL in the absence of dynactin and dynein-dynactin in the absence of NINL (Figure 2A, C). To do this, we varied the concentration of NINL or dynactin between 0.005 and 0.010 μM and then calculated the fraction of dynein bound to NINL or dynactin over time. We ensured that dynein on coverslip was sparse (calculated concentrations of dynein for each kinetic experiment in this study did not exceed 1 pM), meaning that association with dynein does not significantly deplete the solution concentration of dynactin or adaptor. For each experiment, the individual binding curves fit a single exponential, as expected for pseudo-first order binding, and the measured k_obs_ displayed a linear dependence on NINL or dynactin concentrations over the range we explored (Figure S2A-D).

**Figure 2:**
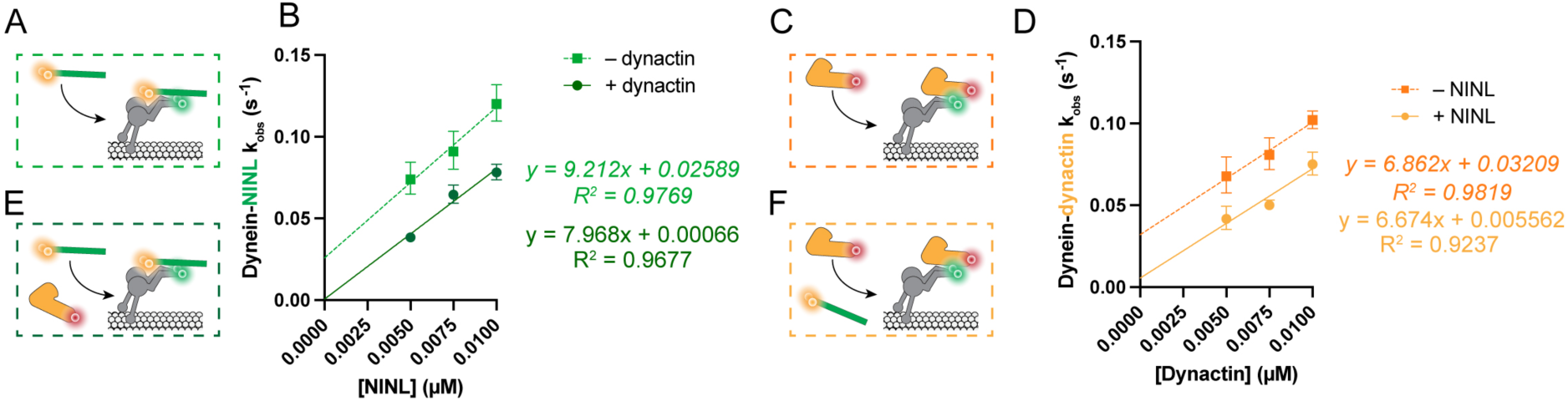
Binding of NINL and dynactin with dynein is not cooperative. **A.** Diagram of NINL association with dynein in the absence of dynactin. **B.** k_obs_ of NINL binding to dynein at varied NINL concentrations in the presence (dark green) or absence (light green) of dynactin with corresponding linear regression equations. The data in the absence of dynactin is normalized for the difference in off rates as described in the text. **C.** Diagram of dynactin association with dynein in the absence of NINL. **D.** k_obs_ of dynactin binding to dynein at varied dynactin concentrations in the presence (yellow) or absence (orange) of NINL with corresponding linear regression equations. The data in the absence of NINL is normalized for the difference in off rates as described in the text. **E.** Diagram of NINL association with dynein in the presence of dynactin. **F.** Diagram of dynactin association with dynein in the presence of NINL.

The measured k_obs_ reflect contributions from both k_on_ and k_off_ because under these conditions k_obs_ = [ligand]*k_on_ + k_off_, where ligand is either NINL or dynactin.^18^ As an internal control, we directly measured k_off_ values for the dynein-NINL and dynein-dynactin at each concentration range tested, anticipating that the measured k_off_ values should be independent of NINL or dynactin concentration. Contrary to our expectations, we observed a linear relationship between NINL and dynactin concentration and the measured k_off_ rates, with higher protein concentrations resulting in decreased off-rates (Figure S2E and F). We reasoned that we may be missing bona fide unbinding events due to the 1.2 second cut-off imposed in our analysis. If k_on_ rates were sufficiently fast, rapid NINL or dynactin unbinding and rebinding could be mistaken for a prolonged binding event. Because k_on_ is concentration dependent, this effect would disproportionally effect experiments with higher protein concentrations. To test this, we repeated the dynein-NINL binding experiment with 0.010 μM NINL, labeling half of the NINL with JF646 and the other with TMR. We observed that the dwell time of dynein colocalized with NINL-JF646 or NINL-TMR was shorter than the measured dwell of dynein colocalized with NINL, irrespective of dye color, which indicates that rapid unbinding and rebinding is occurring (Figure S2G). In fact, by inspecting individual dynein ROIs, we were also able directly observe rapid NINL exchange, which would have been missed if both NINL molecules were labeled with the same dye (Figure S2H). Together, these data indicate that rapid rebinding in our assay can result in missed unbinding events and adds concentration-dependent error to the measured k_obs_ values. To correct our k_obs_ values for missed unbinding events, we first fit the measured k_off_ rates to a line to identify the y-intercept as an approximation of the true k_off_ (Figure S2E and F). For dynein-NINL, this value was 0.0627 sec^-1^ and for dynein-dyactin it was 0.0817 sec^-1^. Next, we added the difference between the true k_off_ and the measured k_off_ at each concentration of NINL or dynactin to the measured k_obs_ (Figure 2B and D). With this correction, the association rate constant for dynein-NINL was 9.21 μM^-1^ sec^-1^ and for dynein-dynactin was 6.86 μM^-1^ sec^-1^.

Next, we set out to measure the kinetics of NINL and dynactin binding to dynein when all three components were present (Figure 2E, F). To perform these experiments, we varied the concentration of either NINL or dynactin while holding the other constant and monitored the fraction of dynein molecules bound to the variable component over time. It is possible that dynactin and adaptor could interact with each other prior to dynein binding, which would increase the complexity of the reaction scheme and our complicate our ability to interpret the measured association rates. To test this possibility, we measured NINL-dynactin binding by a pull-down assay at concentrations 10-fold higher than used in this study and detected no interaction (Figure S1E). Therefore, we did not consider any potential effect of dynactin-adaptor association before dynein binding in these and all following experiments where dynactin and an adaptor were present together.

The resulting binding curves for both dynein-NINL in the presence of dynactin and dynein-dynactin in the presence of NINL fit to single exponentials (Figure S2I and J). Following our previous approach, we next measured the k_off_ rates for dynein-NINL and dynein-dynactin complexes at each concentration tested. Unlike what we observed with the bimolecular binding experiments, the k_off_ values measured when all three components were present did not vary with concentration (Figure S2E and F). Further, the measured k_off_ rates when all components were present were significantly slower than the measured off-rates for either biomolecular experiment (Figure S2E and F). This result indicates that rapid rebinding to dynein did not introduce concentration dependent error to the k_obs_ for binding experiments performed with all three components. We reason that, because the off-rates are significantly slower, the initial complexes formed remain bound for a longer proportion of the experiment and therefore there are fewer rapid rebinding events that erroneously lengthen the apparent complex dwell time.

The k_obs_ for dynein-NINL with dynactin and dynein-dynactin with NINL were linear over the range of concentrations tested and yielded association constants of 7.97 μM^-1^ sec^-1^ for dynein-NINL in the presence of dynactin and 6.67 μM^-1^ sec^-1^ for dynein-dynactin in the presence of NINL (Figure 2B, D). Importantly, these association rate constants are very similar to those measured for dynein-NINL and dynein-dynactin in the absence of the other binding partner, which indicates that under the conditions of the experiments, NINL and dynactin’s association with dynein are likely independent and not cooperative. Moving forward, we will use the average of the association constants measured in the two- and three-component experiments to describe dynein-NINL and dynein-dynactin binding (8.59 μM^-1^ sec^-1^ and 6.77 μM^-1^ sec^-1^, respectively). Together, these results indicate that, for instance, the binding of one component to dynein does not induce a conformational change that enhances the binding rate of the second component.

### A population of activated dynein complexes are very long lived

While association rates were similar for both NINL and dynactin in the bimolecular and trimolecular assembly experiments, the measured off-rates were different. To further probe dynein’s interaction with NINL or dynactin, we first examined the distributions of dwell times of dynein complexes formed with either 0.010 μM NINL or dynactin in the absence of the other component. While there was a major peak for both dynein-NINL and dynein-dynatin dwell times at 5 seconds, both dwell time distributions were complex and fit best to multiple Gaussians (Figure 3A and B). To more easily quantify the complex stabilities from the measured dwell times, we grouped the measured dwells into three categories: short (events that lasted <20 seconds), medium (events that lasted 20-80 seconds), and long (events that lasted >80 seconds). dynein-NINL were more stable with 47% of complexes short-lived, 37% medium-lived, and 16% exceeded 80 seconds (Figure 3C). In contrast, dynein-dynactin bimolecular complexes were less stable, with 69% short-lived, 22% medium-lived, and only 9 % long-lived (Figure 3D). Together, these data indicate that dynein’s interaction with both NINL and dynactin alone are generally short, with dynein-dynactin events being shorter than dynein-NINL. The observed dwell time differences between NINL and dynactin complexes are also consistent with the off-rates measured and described above (Figure S2E and F)

**Figure 3:**
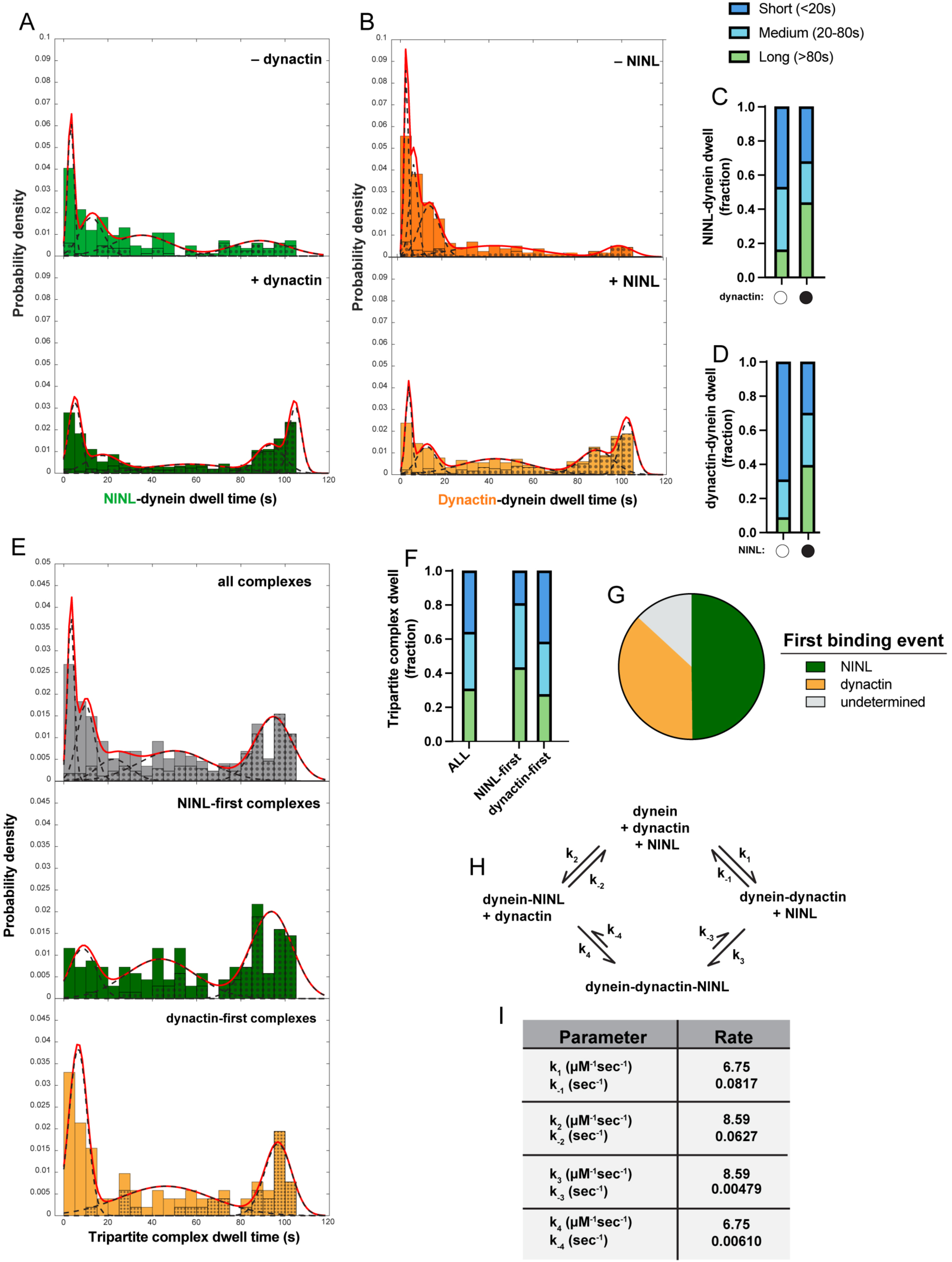
Subpopulations of dynein-dynactin-NINL complexes and more stable complexes form via NINL-first binding events. **A.** Histograms of the dwell time of NINL on dynein in the absence (light green) and presence (dark green) of dynactin. Overall probability density function is shown as a solid red line, with individual component Gaussians as dashed black lines. Black dots show the percentage of the data in each bin that was truncated by the end of the acquisition. **B.** Histograms of the dwell time of dynactin on dynein in the absence (orange) and presence (yellow) of NINL. Overall probability density function is shown as a solid red line, with individual component Gaussians as dashed black lines. Black dots show the percentage of the data in each bin that was truncated by the end of the acquisition. **C, D.** Fraction of events with dwells that fall into short (<20s), medium (20-80s) or long (>80s) timescales. Fraction shown in (C) is calculated from data shown in (A); fraction in (D) is calculated from data shown in (B). **E.** Histograms of the dwell time of dynein-dynactin-NINL complexes. All of the observed complexes are shown in gray, with the complexes that had NINL bind dynein first in green and the complexes that had dynactin bind dynein first in yellow. Overall probability density function is shown as a solid red line, with individual component Gaussians as dashed black lines. Black dot shows the percentage of the data in each bin that was truncated by the end of the acquisition. **F.** Fraction of events with dwells that fall into short (<20s), medium (20-80s) or long (>80s) timescales, as calculated from data shown in (E). **G.** Percentage of complexes that had NINL or dynactin bind dynein first. Undetermined refers to complexes in which NINL and dynactin both bound within one 0.15 s frame. **H.** Reaction scheme for the formation of the activated transport complex with dynactin and NINL. **I.** Kinetic rates for the reaction scheme in (H).

To determine how the presence of the other components alters binding stability of the complexes, we next plotted the dwell time distributions of dynein-NINL and dynein-dynactin obtained from the experiments where both NINL and dynactin were present at 0.010 μM (Figure 3A and B). Like the two component experiments, the dwell time distributions from dynein-NINL and dynein-dynactin complexes formed when all three components were present fit to multiple Gaussians and there was a significant population of short-lived events. However, when all three components were included in the experiment, we observed a dramatic increase in the proportion of long-lived events, with 44% dynein-NINL and 39% dynein-dynactin complexes exceeding 80 seconds (Figure 3C and D). It is also important to note that nearly 50% of total events were terminated by the movie ending and not by dissociation, which means that the measured dwells are an underestimation of actual dwell times. This observation of long-lived complexes is consistent with the slower k_off_ values measured in experiments when all three components are included (Figure S2E, F).

Despite the clear population of long-lived complexes formed when all three components were present, there was also a population of short-lived binding events (32% and 30% for dynein-NINL and dynein-dynactin, respectively) (Figure 3C, D). From the way that we’ve conducted the analysis thus far, it is possible that the short-lived dynein-NINL and dynein-dynactin complexes observed in the three component binding experiments could correspond to events where only dynactin or NINL is bound (i.e. biomolecular binding events) and that the longer-lived species only occurred when all three components were present at one dynein ROI (i.e. tripartite complex formation). To test if this was true, we explicitly extracted dwell times from events only where all three components were colocalized. We observed that dwell times of trimolecular complexes displayed both a short- and long-lived species (Figure 3E, F). This indicates that both low and high stability dynein-dynactin-NINL tripartite complexes can form.

### NINL-first binding events favor more stable complexes

An outstanding question is whether there is a required assembly order for forming the activated transport complex. In other words, *is the binding order of dynactin and adaptors to dynein random, or must one of the components bind first?* The measured association rate constants of NINL-dynein and dynactin-dynein are similar, which means that both binding events occur in similar time frames (Figure 2B and D). Because we conducted the trimolecular experiments with fluorophores on both NINL and dynactin, it was possible to monitor each event to determine which component bound first. Consistent with their similar association rates, we observed that 49% of tripartite complexes form NINL-first binding events and 37% form from dynactin-first binding events. (Figure 3G). NINL and dynactin appear to land simultaneously for 13% of tripartite complexes, which is likely a result of the 150 msec detection limit of our movies. Importantly, the ratio of NINL-first to dynactin-first events (49% / 37% = 1.32) is consistent with what would be predicted from simply considering the ratio of the measured association rate constants (8.59 μM^-1^ sec^-1^ / 6.77 μM^-1^ sec^-1^ = 1.27). Together, these data suggest that dynein’s association with NINL and dynactin is random and simply governed by their respective association rate constants.

Next, we asked *does the order of binding influence the stability of the resultant complex?* To answer this, we compared the dwell times of tripartite complexes that formed via NINL-first binding events to those that formed via dynactin-first binding events. Surprisingly, we found that complexes that formed via NINL binding first were more stable, with 43% of events surviving for greater than 80 seconds and only 19% of complexes being short-lived. In contrast, only 27% of events that formed via dynactin-first binding survived longer than 80 seconds, while 41% were short-lived (Figure 3E, F). We interpret this to mean that stable, long-lived dynein-dynactin-NINL complexes are most likely to form when NINL binds first.

Once a tripartite complex is formed, it could disassemble either by dynactin or NINL dissociating first. Thus, next we asked *does the tripartite complex had a preferred pathway for disassembly?* To address this question, we calculated the off-rates of NINL and dynactin from tripartite complexes. We observed no difference in the measured k_off_ for NINL or dynactin’s dissociation from the tripartite complex (Figure S3A). This means that once formed, NINL and dynactin are equally likely to unbind from the complex.

With the data we have collected thus far, we can now generate and populate a reaction scheme for the formation of the activated transport complex with dynactin and the adaptor NINL (Figure 3H and I). Our observation that either NINL and dynactin can bind first indicates that the binding reaction is not ordered, and that formation of the complex can proceed via both arms of the diagram. Because we observe similar rate constants for dynein’s association with both partners in the bimolecular and trimolecular binding reactions, we can assume that k_1_ = k_4_ and k_2_ = k_3_ (Figure 3H). For these rate constants, we report the average of the association constants measured in the bimolecular and trimolecular experiments (Figure 2B and D). The dissociation rate constants k_-1_ and k_-2_ are calculated from the off-rates measured in the bimolecular experiments (Figure S2E and F), and k_-3_ and k_-4_ are calculated directly from the trimolecular binding experiments (Figure S3A).

### Adaptor identity controls association kinetics and mechanism of activated dynein complex formation

Different adaptors link dynein to different cargos.^4,17^ We hypothesized that different adaptors may support activated transport complex formation with different kinetics. To test this hypothesis, we used the real-time association assay to monitor dynein-dynactin-adaptor complex formation with the adaptor BicD2, which links the dynein transport machinery to Golgi derived vesicles and the Nuclear Pore Complex.^19–23^ Here, we used a well-characterized C-terminal truncation of BicD2 that prevents autoinhibition.^16^ In traditional motility assays performed at equilibrium, both NINL and BicD2 promote robust formation of activated transport complexes that move processively along microtubules with similar velocities.^16,24^ In contrast, we found that in the real-time association assay, NINL and BicD2 behaved quite differently. In the absence of dynactin, there was almost no measurable binding between dynein and 0.010 μM BicD2 (Figure S4A). The few dynein-BicD2 binding events that did occur were short lived, with a measured off-rate of 0.137 sec^-1^ (Figure 4A, B and S4B).

**Figure 4:**
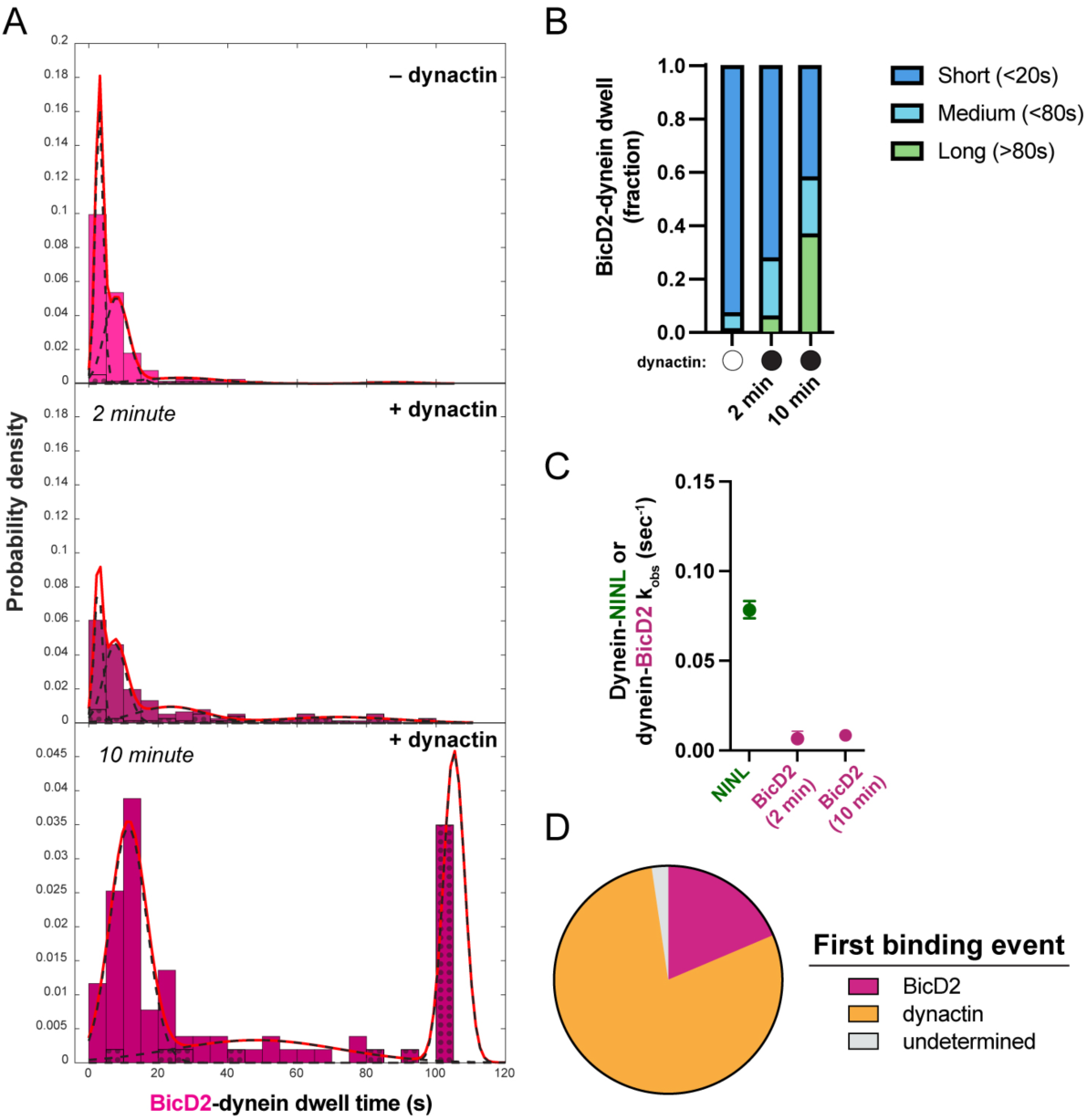
Adaptor identity controls the assembly kinetics of dynein activated transport complexes. **A.** Histograms of the dwell time of BicD2 on dynein in the absence (light pink) and presence (dark pink) of dynactin. Data in the presence of dynactin was collected for 2 min at 0.15 s/frame and at 10 min at 0.75 s/frame. For the 10-minute data any dwell times greater than 110 seconds were randomly given a value between 100-110 and marked as censored for this histogram to better compare to the 2-minute data. Overall probability density function is shown as a solid red line, with individual component Gaussians as dashed black lines. Black dots show the percentage of the data in each bin that was truncated by the end of the acquisition. **B.** Evens with dwells times from (B) that falls into short (<20s), medium (20-80s) or long (>80s) timescales, calculated from data shown in (A). **C.** k_obs_ for dynein-NINL and dynein-BicD2 in the presence of dynactin. NINL data repeated from Figure 2B. **D.** Percentage of complexes that had BicD2 or dynactin bind dynein first.

Next, we monitored dynein-BicD2 association in the presence of 0.010 μM dynactin. Although we observed modest association of BicD2 with dynein in these conditions, the fraction of dynein molecules bound by the end of data acquisition never exceeded ∼25% (Figure S4C). For comparison, ∼60% of dynein molecules were bound to NINL under identical conditions (Figure S2A). Similarly, dynein-BicD2 complexes that formed in the presence of dynactin were less stable, with 72% of complexes being short-lived, 22 % medium-lived, and only 6% long-lived (Figure 4A and B). This is contrast to dynein-NINL complexes, in which over 44% percent had dwells longer than 80 seconds in the presence of 0.010 μM dynactin (Figure 3A and C). A similar trend was observed in the distributions of dwell times for dynein-dynactin-BicD2 tripartite complexes (Figure S4D and E). Consistent with these observations, the off-rate of dynein-Bicd2 complexes in the presence of dynactin (0.03799 sec^-1^) was much faster than the equivalent measurement with NINL (0.00803 sec^1^) (Figure S2E, S4B). Because BicD2 displayed such poor binding to dynein at concentrations amenable to single-molecule analysis, we were unable to determine a dynein-BicD2 association rate constant. However, the k_obs_ for dynein-BicD2 binding in the presence of dynactin is an order of magnitude less than the k_obs_ for NINL in equivalent conditions (Figure 4C).

These data suggest that BicD2 binds slower and forms less long-lived complexes with dynein than NINL. This result also suggests that the 2-minute duration of our association experiment may be too short to observe appreciable BicD2 binding. Thus, we repeated the real time association assay with BicD2 and dynactin but allowed binding to occur over a ten-minute period. To minimize bleaching, we imaged every 750 msec rather than every 150 msec. Consistent with slower binding, performing the experiment over a period of ten minutes allowed us to observe an appreciable fraction of the dynein ROIs become populated with BicD2 molecules (Figure S4F). As expected, the k_obs_ determined for dynein-BicD2 in the 10- and 2- minute experiments were the same (Figure 4C). During this experiment, we also monitored dynactin binding and observed much faster association kinetics for dynactin than for BicD2 (Figure S4F). The difference in dynein-dynactin and dynein-BicD2 association rates means that most complexes should form via dynactin-first binding, which we confirmed via directly monitoring the order of assembly for each tripartite complex (Figure 4D). We were also able to observe a significant increase in the number of dynein-BicD2 and dynein-dynactin-BicD2 tripartite complexes that were long-lived, with 37% of dynein-BicD2 complexes and 35% of tripartite complexes surviving longer than 80 seconds (Figure 4A, B, S4D-H). Together these data show that because BicD2 binds dynein with significantly slower binding kinetics than NINL, dynein-dynactin-BicD2 complexes form much more slowly than complexes with NINL and assemble primarily via dynactin binding first. However, once formed, the dynein-dynactin-BicD2 and dynein-dynactin-NINL complexes have comparable stabilities.

Because NINL and BicD2 behave similarly in motility assays, these results were very surprising^16^. However, it should be noted that the motility studies assessed dynein activation after a 20-minute incubation of dynein, dynactin, and adaptor.^16^ We reasoned that, by performing “kinetic motility” experiments in which the pre-incubation time of dynein, dynactin, and adaptor is varied, we should be able to directly observe the effects of assembly kinetics on dynein motility. To perform these experiments, we mixed dynein, dynactin, and adaptor for between 1 minute (the fastest we could mix the proteins and image) and 20 minutes; we then added ATP, imaged motility on Taxol-stabilized microtubules with TIRF microscopy. We assessed complex formation by determining the processive complex landing rate from the motility movies (Figure 5A). Consistent with fast association, NINL’s ability to generate activated transport complexes plateaued by the first time point (1 minute), suggesting rapid complex formation (Figure 5B, C). In contrast, we observed results with BicD2 that were consistent with a dramatically slower association rate. With increasing incubation times, there was a clear increase in the amount of processive complexes formed, as measured by the landing rate (Figure 5B, C). By 20 minutes, BicD2 and NINL experiments had the same processive landing rate, which explains how the differences between the association kinetics of these adaptors previously eluded us (Figure 5B, C). Altogether, these data show that different adaptors can facilitate the assembly of activated complex formation with different kinetics. These differences also support the need to examine the dynamic process of dynein activation in pre-equilibrium conditions, as these adaptors don’t diverge from each other dramatically once they have formed activated transport complexes.

**Figure 5:**
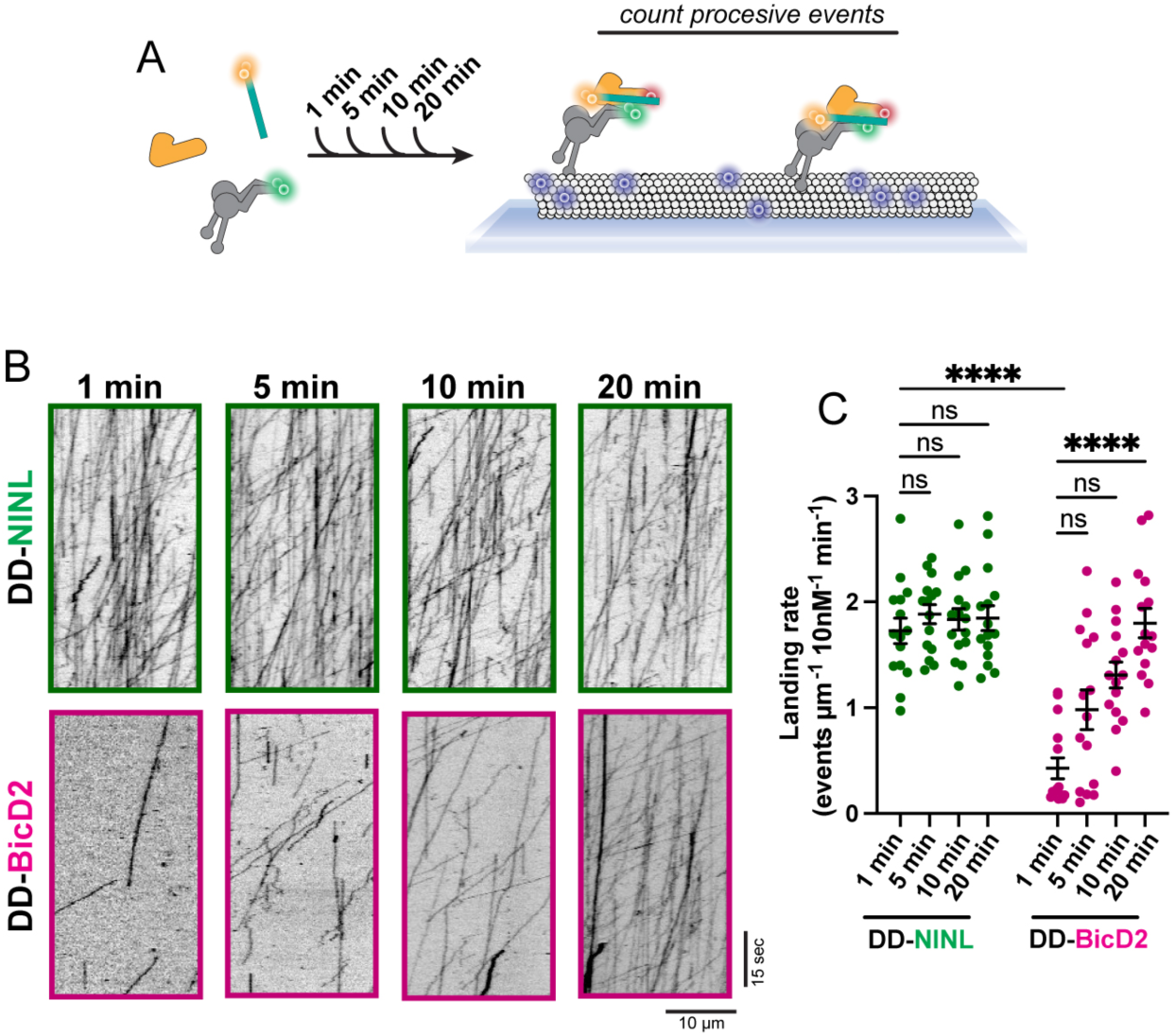
BicD2 activates dynein motility more slowly than NINL. **A.** Diagram of a kinetic motility experiment in which dynein, dynactin and adaptor are incubated for 1, 5, 10, or 20 minutes before imaging dynein motility. **B.** Example kymographs from the kinetic motility assay. **C.** Landing rate of motile dynein-dynactin-NINL (green) or dynein-dynactin-BicD2 (pink) complexes. n = 15 microtubules. Statistical testing was performed with a Kruskal-Wallis test with Dunn’s multiple comparisons test. ns p >0.05, **** p <0.0001.

## Discussion

We set out determine how dynein, dynactin, and cargo adaptors assemble into the activated transport complex. To accomplish this, we developed a single-molecule, TIRF-based assay that allowed us to measure the association kinetics of complex formation and visualize resultant complex stability. To our knowledge, this work is the first systematic effort to describe the dynamic, molecular processes that lead to dynein activation.

Using dynactin and the cargo adaptor NINL, we determined the mechanism by which the activated transport complex forms. Our work showed that the pairwise association of dynein with either dynactin or NINL are independent from each other. In other words, NINL and dynactin interact with free dynein at the same rate that they interact with dynein bound to the other partner, and complex formation can occur via dynactin or NINL binding first. Additionally, the association rate constants of dynein-dynactin and dynein-NINL binding are similar to one another, meaning that at similar free concentrations, complex assembly will occur roughly equally by the two independent pathways (Figure 3H, I).

Despite both pathways being available and their association rates being kinetically comparable, we observed that complexes formed via NINL-first events were more likely to be long-lived than complexes formed via dynactin-first events. Why might dynactin-first complexes be less likely to yield stable complexes? We reason that dynactin binding first can result in an “off-pathway” intermediate that has a greater probability of generating a complex with reduced stability. The contacts between dynein, dynactin, and an adaptor in the assembled activated transport complex are extensive and multivalent.^7,8,11^ Adaptors interact with dynein light intermediate chain via a domain at their N-terminus and interact with multiple sites on dynein heavy chain via an extended coiled coil (Figure 1A).^7,8,11,25–27^ Adaptors also bind to dynactin’s Arp filament via their coiled-coil and dynactin’s pointed end complex via a short motif within the coiled-coil, called a spindly motif.^28^ Dynein and dynactin also have multiple points of contact. Dynein intermediate chain binds to a dynactin’s p150 subunit and dynein heavy chain makes multiple sites of contact with dynactin subunits along the filament (Figure 1A).^8,29,30^ In the final active structure, the adaptor appears sandwiched between dynactin’s Arp filament and the tails of the dynein heavy chain. We speculate that dynactin-first events may be prematurely “zipped-up” such that some of contacts between dynein heavy chain and dynactin form and disfavor or preclude adaptor insertion. In contrast, binding of the adaptor to dynein may allow normal dynein-dynactin contacts to form, resulting in a higher probability of stable assembly when adaptors bind first. More work is required to understand complex assembly from a structural perspective, to determine which protein-protein contacts drive association, and to identify the structure of assembly intermediates.

We also found that the adaptors NINL and BicD2 bind dynein with dramatically different association kinetics, which causes BicD2 to form activated transport complexes an order of magnitude slower than NINL. This also means that dynein-dynactin-BicD2 complexes assemble primarily via the dynactin-first pathway. Despite slower association kinetics, dynein-dynactin-BicD2 complexes can form structures that are stable and long-lived, as assessed by complex dwell time. We were also able to determine how differences in assembly kinetics manifested in traditional motility experiment read-outs of dynein activation and found that dynein-dynactin-BicD2 complexes required significantly longer incubation times to yield activated transport complexes. These findings underscore how important it is to consider the kinetics of dynein binding to its regulatory partners when conducting traditional motility experiments.

The difference in assembly kinetics for complexes formed with NINL and complexes formed with BicD2 has potentially broad and significant implications. Dynein is tethered to its myriad cargos by one of nearly 20 different adaptors.^4^ It is not clear how dynein distinguishes between different cargo adaptors to traffic the appropriate cargo. Our work suggests that distinct association rates between dynein and different adaptors could be one mechanism by which dynein discriminates between adaptors to achieve trafficking specificity.

It is also possible that the differences in association rates between NINL and BicD2 have evolved as such to offset other kinetic processes that govern dynein’s ability to traffic cargo. For example, both adaptor constructs used in this study are truncated, thus relieving potential autoinhibition. BicD2 is known to be autoinhibited and can’t associate with dynein when its C-terminal tail is folded back to interact with the N-terminal coiled-coil.^15,20^ To facilitate switching between the folded, autoinhibited to open conformation, BicD2 binds to cargo-localized regulators, like Nup358 or Rab6.^15,31–33^ These same proteins promote BicD2’s cargo localization, which means that BicD2 on cargo is likely in the open state that is competent to bind dynein.

One could imagine that the slow association kinetics of dynein-BicD2 binding is essential for preventing excessive dynein activation on cargo that is laden with open BicD2. It is also possible that the differences in dynein-adaptor association kinetics have evolved to compensate for differences in adaptor concentration or accessibility. During interphase, NINL localizes primarily to the centrosome.^34^ It is unclear what the structure of NINL is at the centrosome and thus, it is unclear whether dynein has access to all NINL molecules. Given the density and layered ultrastructure of the centrosome, it is possible that NINL is largely inaccessible to dynein and that rapid association kinetics ensures that the few accessible NINL molecules can drive activated transport complex formation.^35^

By developing a novel in vitro assembly assay, we show that different adaptors bind to dynein with different kinetics, resulting in different assembly pathways for dynein-dynactin-adaptor complex formation. The present study sets the stage for a more complete accounting of how differences in adaptor association kinetics are harnessed by the cell to promote dynein’s multiple functions. An important next step will be to determine the association rate constants for dynein’s interaction with all other adaptors to fully map the kinetic parameters that govern dynein activation on different types of cargos.

## Methods

### Protein expression and purification

Dynein was expressed in Sf9 cells as described.^24,36^ Briefly, human dynein genes in pACEBac1 plasmid was transformed into DH10EmBacY cells via 42°C heat shock for 15 seconds followed by a 6 hour outgrowth at 37°C in S.O.C. media (Thermo Fisher Scientific) with shaking at 220 rpm. Cells were then plated on LB-agar containing kanamycin (50 μg/mL), gentamicin (7 μg/mL), tetracycline (10 μg/mL), BluoGal (100 μg/mL) and IPTG (40 μg/mL). After 48-72 hours growth, white colonies were selected. Colonies were tested via PCR for the presence of all dynein genes. Selected colonies were grown 37°C with shaking at 220 rpm in LB media with kanamycin (50 μg/mL), gentamicin (7 μg/mL) and tetracycline (10 μg/mL). After 14-16 hours of growth, bacmid DNA was then extracted from cultures with isopropanol extraction. 1 - 2 µg of purified bacmid was transfected into 1 x 10^6^ Sf9 cells with FuGene HD transfection reagent, according to manufacturer’s instruction (Promega). Transfected cells were incubated in a 6 well dish at 27°C without agitation for 3 days. 1mL of supernatant (designated V0) from the transfected cells was collected via centrifugation (1000 x g, 5 min, 4°C) and used to transfect of 50 x 10^6^ SF9 cells/mL, in a final volume of 50 mL. Cells were agitated at 105 rpm at 27°C for 3 days, before the supernatant (designated V1) was harvested via centrifugation. To express protein, 4 mL of V1 was used to transfect 400 x 10^6^ SF9 cells/mL in a final volume of 400 mL and agitated at 105 rpm at 27°C for 3 days. Cells were harvested via centrifugation, the pellet was washed with 10 mL of cold PBS, then frozen in liquid nitrogen and stored at −80°C until needed for protein purification.

Purification of dynein was conducted as described.^24,36^ All steps are performed at 4°C unless otherwise noted. Briefly, dynein pellets were thawed on ice and resuspended in 40 mL per pellet of dynein-lysis buffer (50 mM HEPES (pH 7.4), 100 mM sodium chloride, 1 mM DTT, 0.1 mM Mg-ATP, 0.5 mM Pefabloc, 10% (v/v) glycerol) supplemented with cOmplete EDTA-free protease inhibitor cocktail (Roche). Lysis was accomplished via douncing and the lysate was clarified via centrifugation (183,960 x g, 88 min, 4°C) in a Type 70Ti rotor (Beckman). 2 mL of IgG Sepharose 6 Fast Flow beads (Cytiva) were incubated with the clarified supernatant for 4 hours with rotation. Beads were collected in a gravity column and washed with at least 200 mL of dynein-lysis buffer and 300 mL of TEV buffer (50 mM Tris–HCl (pH 8.0), 250 mM potassium acetate, 2 mM magnesium acetate, 1 mM EGTA, 1 mM DTT, 0.1 mM Mg-ATP, 10% (v/v) glycerol). For fluorescent labeling, dynein-bound beads were mixed with 5 µM SNAP-Cell-TMR, SNAP-AlexaFluor-647, or SNAP-Alexa-488 (New England Biolabs) for 10 min at RT. Beads were then washed with 300 mL of TEV-buffer, then resuspended in 15 mL TEV buffer supplemented with 0.5 mM Pefabloc and up to 0.2 mg/mL TEV protease and incubated overnight with rotation. Cleaved proteins were separated from the beads with a gravity column and concentrated to 500 µl with a 100K MWCO concentrator (EMD Millipore). Dynein was then injected onto to TSKgel G4000SWXL column (Tosoh) equilibrated with GF150 buffer 25 mM HEPES (pH 7.4), 150 mM KCl, 1 mM MgCl_2_, 1 mM DTT) supplemented with 0.1 mM Mg-ATP. Column was run at 0.75 mL/min, peak fractions were collected, buffer exchanged into GF150 buffer supplemented with 0.1 mg/mL ATP and 10% glycerol, concentrated to 0.1 – 0.5 mg/mL, flash frozen in small aliquots, and stored at −80°C.

Dynactin was purified from HEK293T cells stably expressing p62-Halo-3xFLAG as described.^17^ Briefly, frozen pellets generated from 160 x 15 cm plates of ∼80% confluent cells were resuspended in 80 mL of of dynactin-lysis buffer (30 mM HEPES (pH 7.4), 50 mM potassium acetate, 2 mM magnesium acetate, 1 mM EGTA, 1 mM DTT, 10% (v/v) glycerol) supplemented with 0.5 mM Mg-ATP, 0.2% Triton X-100, and 1x cOmplete EDTA-free protease inhibitor cocktail tablets (Roche)). The lysate was clarified via centrifugation (66,000 x g, 30 min, 4°C) in a Type 70 Ti rotor (Beckman), before being incubated overnight at 4°C with 1.5 mL of anti-FLAG M2 affinity gel (Sigma-Aldrich) with rotation. Beads were collected with a gravity column, then washed with at least 50 mL of wash buffer (Dynactin-lysis buffer supplemented with 0.1 mM Mg-ATP, 0.5 mM Pefabloc and 0.02% Triton X-100), 100 mL of wash buffer supplemented with 250 mM potassium acetate, and then washed again with 100 mL of wash buffer. To elute dynactin, beads were incubated with 1 mL elution buffer (wash buffer with 2 mg/mL of 3xFlag peptide).

Dynactin was then diluted to 2 mL in Buffer A (50 mM Tris-HCl (pH 8.0), 2 mM magnesium acetate, 1 mM EGTA, and 1 mM DTT) and loaded onto a MonoQ 5/50 GL column (Cytiva) at 1 mL/min. A linear gradient from 35 – 100% Buffer B (50 mM Tris-HCl (pH 8.0), 2 mM magnesium acetate, 1 mM EGTA, 1 mM DTT, 1 M potassium acetate) was run over 26 column volumes. Fractions of pure dynactin that eluted between 75 – 80% Buffer B were collected, pooled, and buffer exchanged into GF150 buffer supplemented with 10% glycerol, concentrated to 0.02-0.1 mg/mL, flash frozen in small aliquots, then stored at −80°C.

NINL and BicD2 constructs containing N-terminal HaloTags were expressed and purified as described.^16,24^ Constructs were transformed into BL-21[DE3] cells (New England Biolabs) cells and cell were grown at 37°C with shaking until they reached to an optical density at 600 nm of 0.4-0.6. Then the temperature was reduced to 18°C and protein expression was induced with 0.1 mM IPTG for 16 hours. Pellets were harvested via centrifugation, frozen, and stored at - 80°C until needed. All purification steps were performed at 4°C, unless indicated otherwise. To purify either adaptor, pellets harvested from 1.5 L were thawed in 40 mL of adaptor-lysis buffer (30 mM HEPES pH 7.4, 50 mM potassium acetate, 2 mM magnesium acetate, 1 mM EGTA, 1 mM DTT and 0.5 mM Pefabloc, 10% (v/v) glycerol) supplemented with 1x cOmplete EDTA-free protease inhibitor cocktail tablets (Roche) and 1 mg/mL lysozyme. To lyse, cell slurry was sonicated then clarified via centrifugation (66,000g for 30 min) in Type 70 Ti rotor (Beckman). The supernatant was mixed with 2 mL of IgG Sepharose 6 Fast Flow beads (Cytiva) for 2 h with rotation before being washed with adaptor-lysis buffer supplemented with 150 mM potassium acetate and 50 mL of cleavage buffer (50 mM Tris-HCl pH 8.0, 150 mM potassium acetate, 2 mM magnesium acetate, 1 mM EGTA, 1 mM DTT, 0.5 mM Pefabloc and 10% (v/v) glycerol). The washed beads were then resuspended in 15 mL of cleavage buffer supplemented with 0.2 mg/mL TEV protease and incubated overnight with rotation. The supernatant containing the adaptor proteins was collected, concentrated to 1 mL with a 30 kDa MWCO concentrator (EMD Millipore), filtered, diluted to 2 mL in Buffer A (30 mM HEPES pH 7.4, 50 mM potassium acetate, 2 mM magnesium acetate, 1 mM EGTA, 10% (v/v) glycerol and 1 mM DTT) and injected into a MonoQ 5/50 GL column (Cytiva) at 1 m/min. A linear gradient was run from 0 – 100% Buffer B (30 mM HEPES pH 7.4, 1 M potassium acetate, 2 mM magnesium acetate, 1 mM EGTA, 10% (v/v) glycerol and 1 mM DTT) over 26 column volumes. Peak fractions containing Halo-tagged adaptors were collected, concentrated to 0.2 mL, diluted to 0.5 mL in GF150 buffer and injected onto a Superose 6 Increase 10/300 GL column (Cytiva) equilibrated with GF150 buffer. Column was run at 0.5 mL/min, peak fractions were collected, and buffer exchanged into GF150 buffer supplemented with 10% glycerol. Proteins were flash frozen in small aliquots before being stored at −80°C.

### Single molecule TIRF microscopy

All single-molecule imaging experiments were performed on an inverted microscope (Nikon, Ti2-E Eclipse) with an 100X, 1.49 N.A. oil immersion objective (Nikon, Apo) equipped with a LUNF-XL laser launch (Nikon) containing 405 nm, 488 nm, 561 nm, and 640 nm lasers. The excitation path was filtered through the appropriate quad bandpass filter cube (Chroma). The emission path was split with the appropriate dichroic mirror and filtered with appropriate emission filters (Chroma) using a W-View GEMINI image splitter (Hamamatsu). All emission signals were detected on an iXon Ultra 897 electron-multiplying CCD camera (Andor Technology). Microscopy controls and image acquisition were controlled with NIS Elements Advanced Research software (Nikon).

#### Motility assay and analysis

Motility experiments were performed in flow chambers assembled as described using No. 1-1/2 coverslips (Corning) functionalized with PEG-biotin.^24,37^ 20 µM bovine tubulin (with ∼10% biotin-tubulin and ∼10% Alexa-488 tubulin) was polymerized for 30 min at 37°C before being stabilized with 20 µM taxol. Flow chambers were assembled by first flowing in 1 mg/mL streptavidin in assay buffer (30 mM HEPES [pH 7.4], 2 mM magnesium acetate, 1 mM EGTA, 10% glycerol, 1 mM DTT, and 20 µM taxol) and incubating for 3 min. Next chambers were washed twice with 20 µL of assay buffer before a fresh dilution of taxol stabilized microtubules (final concentrations ∼0.1 – 0.4 µM) was flowed in and incubated for 3 minutes. Chambers were then washed twice with assay buffer supplemented with 1 mg/mL casein. To assemble dynein-dynactin-adaptor complexes, dynein (15 nM), dynactin, and the adaptor were mixed at a 1:2:10 molar ratio and incubated for between 0 and 19 minutes. Immediately before imaging, the samples were mixed with assay buffer supplemented with 20 µM Taxol, 1 mg/mL casein, 71.5 mM β-mercaptoethanol, 0.05 mg/mL glucose catalase, 1.2 mg/mL glucose oxidase, 0.4% glucose, and 2.5 mM Mg-ATP. Samples were then introduced into the flow chamber and imaged immediately. Microtubules were imaged first by taking a single frame snapshot. Dynein (labeled with JF646) and adaptors (labeled with TMR) were imaged every 300 msec for 2 minutes. At the end of image acquisition, the microtubule channel was imaged once more to assess stage drift and movies showing drift were not used. Kymographs were generated from movies using ImageJ macros as described.^38^ Dynein landing rate was determined per microtubule and calculated by dividing the total number of processive events analyzed for that microtubule by the length of the microtubule, divided by the concentration of adaptor used in that experimental condition, divided by the length of the movie in minutes. Only runs longer than 8 pixels were included in the analysis. Bright protein aggregates, which were defined as molecules 4 times brighter than the background, were also excluded. All statistical analysis was performed in Prism10 (GraphPad).

#### Association assay

Association experiments were performed in custom built flow chambers with ports on either end to allow sample to be introduced during image acquisition. To make these flow chambers, 1.2 mm diameter holes were drilled into either side of a 1 mm glass slide (Fisher, 12-550-15). Flow chambers were constructed as above, with double sided tape separating chambers with a hole on either end. The open ends of the chambers were sealed with epoxy (Devcon). 200 µL pipette tips were inserted into the holes and affixed with epoxy, after which 1.2 mm diameter (0.76 mm inner diameter) tubing was inserted into one of the pipette tips and affixed inside the pipette tip with epoxy. A 20-gauge needle on a 1 mL syringe was inserted into the tubing. To introduce sample into the flow chamber, the sample was pipetted into the pipette tip on the opposite end and the syringe was pulled back until all of the liquid flowed into the chamber.

To assemble chambers for the association experiments, 1 mg/mL streptavidin in assay buffer was incubated for 3 minutes before being washed out with assay buffer. Next, taxol stabilized Alexa-405 labeled microtubules were incubated for 3 min, before being washed out with assay buffer supplemented with 1 mg/mL casein. Dynein labeled with Alexa-488 was diluted to 25 – 100 pM in assay buffer with 1 mg/ml casein, supplemented with 0.025 U/mL apyrase, and then was flowed in and incubated for 10 min. The concentration of dynein in the chamber was calculated by taking the number of dynein spots in a field of view, dividing by Avogadro’s number to get moles and then by the volume of a field of view to get molarity. The volume is calculated from the area of our field of view (81.92 x 40.96 µM) and the height of the double-sided tape we use to make chambers (76.2 µm) which gives 256 pL. A field of view with 150 dynein spots therefore has a dynein concentration of about 1 pM. The fields of view used ranged from 15-150 dynein spots. To monitor association, first a single snapshot of the dynein and microtubule channels were taken before the movie acquisition was started. Next, while imaging the appropriate channels, mixtures of dynactin and/or adaptors labeled with either TMR or JF646 (final concentrations between 5 – 10 nM) were introduced to the chamber. Movies were collected for 2 minutes total, with a frame rate of 150 msec/frame. For longer imaging performed with BicD2, movies were collected for 10 minutes with a frame rate of 750 msec/frame. In both cases, exposure times for each laser were 40 msec. At the end of the movie, another snapshot of the dynein and microtubule channels were acquired to assess stage movement and to ensure dynein molecules were still present.

#### Association assay considerations and analysis

Dynein molecules that were present in the first and last frames and with an were set as regions of interest (ROIs) and circled with a diameter of 6 pixels (0.942 µm). ROIs were assessed manually and those that contained bright puncta indicative of aggregates were not considered. Next, movies of the dynactin and adaptor channels were processed with a rolling ball background subtraction using a 3 pixel radius. Next, traces of the intensity of dynactin or adaptor over time at dynein ROIs were generated and used for analysis. Using custom matlab scripts, we extracted kinetic parameters from each trace.

These parameters include association time, event dwell time, and fraction ROIs bound to dynactin and/or adaptor (Figure 1C). Binding events were defined as positive deviation from baseline intensity, which was set individually for dynactin and adaptor in each dataset. All ROIs were manually checked to determine if a second dynactin or adaptor molecule landed. Dynein ROIs that ever received a second, concurrent binding event were omitted from the analysis in case the ROIs contained two proximal dynein molecules. This occurred for 33% of the ROIs. While we acknowledge that it is possible for two adaptors to bind one dynein, we chose to only consider complexes with a single adaptor (see extended discussion below).

Another possible explanation for multiple dynactin or adaptor molecules appearing to localize simultaneously with one dynein ROI is interactions between dynactin or NINL and the microtubule. To test the contribution of binding to microtubules, we flowed dynactin or NINL into chambers without dynein and found that the landing rate was 0.0053 events/um/min/nM dynactin and 0.0041 events/um/min/nM NINL. Based on the size of our ROIs and this observed landing rate, we would expect about 9.9% of our ROIs to erroneously include a dynactin event and 7.7% to include a NINL event. We conclude that two dynactin or adaptor binding events were likely caused by both two dynein molecules bound close together and interactions of dynactin and adaptor with the microtubule or the antibody. Because only 2% of RO received neither dynactin or NINL, deleting ROIs that received two signals limited the contribution of nonspecific binding.

To calculate the photostability of the fluorophores used in the association assay, we nonspecifically attached labeled dynactin and NINL to a coverslip and analyzed the fluorescent spots with above framework. 32.7% of TMR dyes and 15.2% of JF646 dyes photobleached by the end of the imaging period. Each fluorophore also blinked in our experimental conditions, which makes it potentially challenging to differentiate an unbinding and rebinding event from a dye blink. By plotting a frequency distribution of the time that each blinking event remained in the dark state for each dye, we determined that a cut-off of 1.2 seconds (8 frames in the experiment) would result in 68% (for TMR) and 84% (for JF646) not experiencing a photo event during the 2-minute imaging time. We used this 8-frame persistence cutoff in all 2 min experimental conditions, meaning that all events had to last longer than 1.2 seconds to be considered a bona fide binding or unbinding event. For 10-minute experiments, we adjusted the cutoff to be 4 frames (equivalent to 3 seconds) and applied it in the same way. To ensure that bleaching rates and blinking frequency differences between each dye did not interfere with our analysis, we collected at least one replicate of all experiments with the fluorophore on dynactin and the adaptor swapped.

Association of dynactin or adaptor with dynein was assessed by the fraction of dynein ROIs bound as a function of time.^18^ k_obs_ was determined from each individual binding curve in Prism10 (GraphPad) by fitting to the following equation: 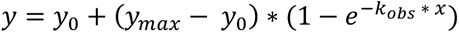. Here, y_0_ is the initial counts and y_max_ is the final counts. Each k_obs_ value was plotted as a function of concentration and fit to a linear regression to determine association rate constants. Off-rates of dynactin and adaptor were calculated by dividing the total number of observed unbinding events by the summed dwell time of every binding event. This approach, which assumes dissociation events are history independent, takes into account binding events that are terminated by the end of the movie rather than an observed dissociation. Off-rates from the complex were calculated in the same way, by taking the number of unbinding events for either dynactin or adaptor and dividing by the summed dwell time of the tripartite complex. Dwell time histograms were generated in Matlab. The number of Gaussians that best fit the data was determined using Bayesian information criterion. The maximum number of Gaussians tested was 6.

#### Experimental limitations and caveats

The association experiments we have designed explicitly monitor the association of one dynein dimer with one dynactin complex and one dimeric adaptor. It is important to emphasize that activated transport complexes often contain two dynein dimers and can also contain two adaptor dimers.^7,8^ To our knowledge, this work represents the first effort towards determining the order of events that precede complex formation, therefore it is not clear if a dynein dimer must bind dynactin and a single adaptor dimer before the second dynein and adaptor are recruited or if these binding events can occur randomly. Determining the sequence of assembly of these higher-order complexes in the future would require, for instance, immobilizing an adaptor and monitoring binding of dynein, dynactin, and an additional adaptor molecule.

### Equilibrium binding experiments

The binding of dynein, dynactin and NINL was measured in solution by coupling 100nM NINL to 35 µL of NEBExpress^®^ Ni-NTA Magnetic Beads (NEB) in the case of NINL-dynactin binding or 100nM dynein to 25 µL SNAP-Capture Magnetic Beads (NEB) in 2 mL Protein Lo Bind Tubes (Eppendorf) using the following protocol. Beads were washed twice with 1 mL of GF150 without ATP supplemented with 10% glycerol and 0.1% NP40. NINL or Dynein was diluted in this buffer to 100 nM. 25 µL of diluted protein was added to the beads and gently shaken for one hour. 20 µL of supernatant were then analyzed via SDS-PAGE to confirm complete depletion of the protein by the beads. The protein-conjugated beads were washed once with 1 mL GF150 with 10% glycerol and 0.1% NP40 and once with 1mL of binding buffer (28.75 mM HEPES [pH 7.4], 1.5 mM magnesium acetate, 0.75 mM EGTA, 0.25mM MgCl_2_, 37.5mM KCl, 10% glycerol, 1 mM DTT, 1 mg/mL casein, 0.1% NP40, 1mM ADP). 5 nM dynactin or NINL was diluted such that the final buffer composition was binding buffer. 25 µL of the dynactin or NINL mixture was added to the beads and gently agitated for 45 minutes. After incubation 20 µL of the supernatant was removed, and 6.67 µL of NuPAGE® LDS Sample Buffer (4X) and 1.33 µL of Beta-mercaptoethanol was added to each. The samples were boiled for 5 minutes before running on a 4-12% NuPAGE Bis-Tris gel. Depletion was determined using densitometry in ImageJ.

## Contributions

JPG and MED conceived of and planned the project. JPG developed the assay. JPG, MED, and WOH established analysis procedures. JPG and SRL performed experiments and analyzed data. MED wrote the manuscript with input from all authors.

## Acknowledgements

The authors thank Drs. Michael Cianfrocco, David Sept, Patrick O’Brien, Kristen Verhey, and Randy Stockbridge as well as members of the DeSantis, Cianfrocco, Verhey, Ohi, and Sept labs for helpful discussions and/or feedback on the manuscript. This work was supported by NIH-R35GM146739 and NSF-2142670 (to MED) and NIH-R35GM139568 (to WOH).

---

**Supplement to Figure 1:**
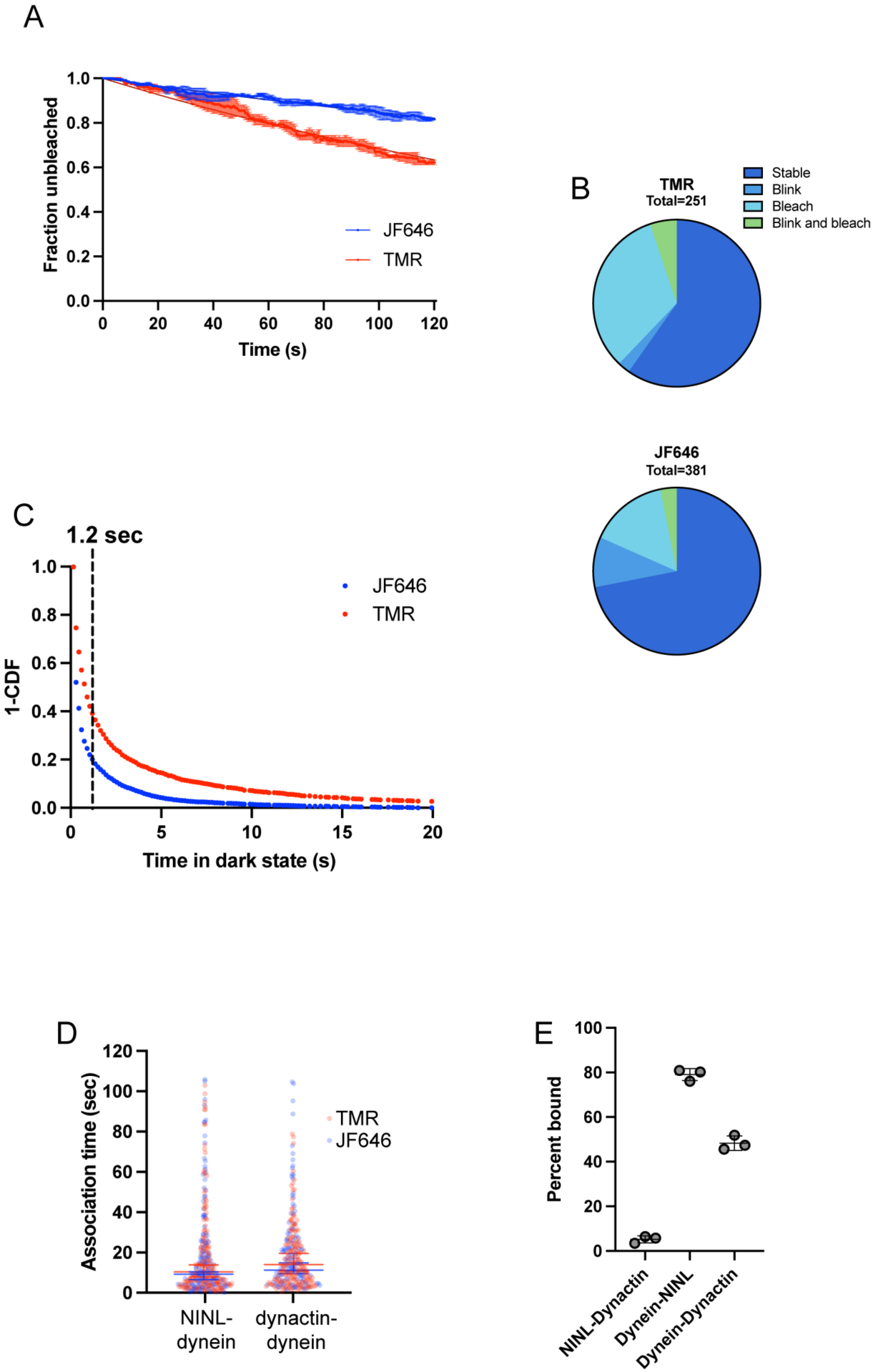
**A.** Fraction of unbleached fluorophores over time for a 120 sec movie. To generate these data, labelled protein was nonspecifically attached to a glass coverslip and imaged under conditions identical to those used in the association assay. The lines represent single exponential fits, with a tau of 356.4 sec (for JF646) and 263.5 sec (for TMR). Error bars are SEM; n = 2. **B.** Percentage of fluorophores that experience a photoevent during the bleaching experiment (in S1A), given a 1.2 sec event cutoff. **C.** 1-Cumulative distribution function of the times that JF646 and TMR dyes remained in the dark state during the bleaching experiment. The 1.2 second cutoff applied in the analysis is indicated with a dashed line. **D.** Time to first association event for NINL-dynein and dynactin-dynein with different fluorophores. Events in which the protein was labelled with TMR are shown in red and events in which the protein was labelled with JF646 are in blue. Error bars are median and interquartile range and are colored by fluorophore. **E.** Percent of dynactin bound to NINL-conjugated beads (NINL-dynactin) and percent of NINL or dynactin bound to dynein-conjugated beads (dynein-NINL and dynein-dynactin).

**Supplement to Figure 2:**
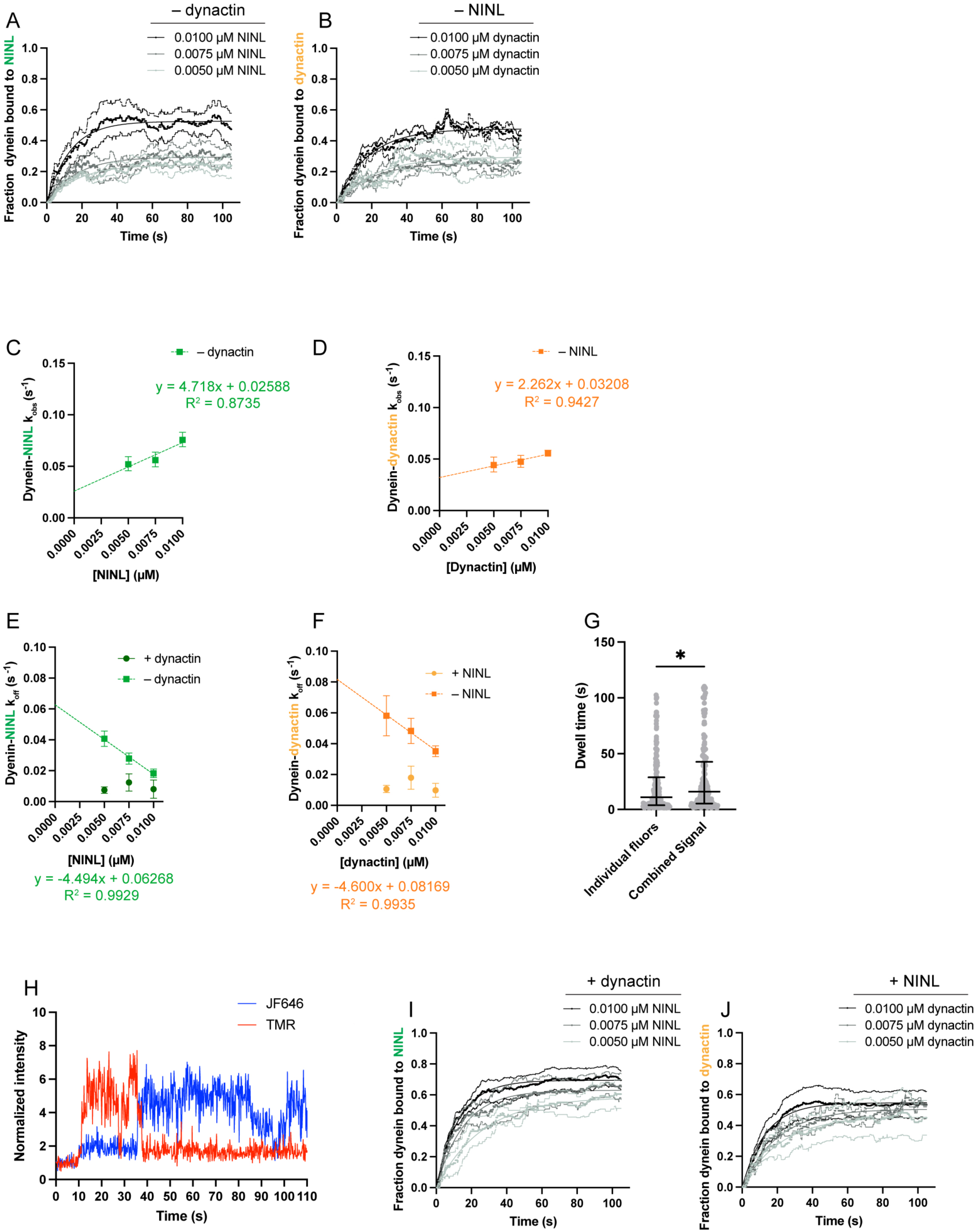
**A.** Fraction of dynein bound to NINL at varied NINL concentrations in the absence of dynactin. Curves are fit to a single exponential. n = 3 for each condition. **B.** Fraction of dynein bound to dynactin at varied dynactin concentrations in the absence of NINL. Curves are fit to a single exponential. n = 3 for each condition. **C.** k_obs_ of NINL binding to dynein at varied NINL concentrations in the absence of dynactin with corresponding linear regression equations. This data is not normalized by the off rate. **D.** k_obs_ of dynactin binding to dynein at varied dynactin concentrations in the absence of NINL with corresponding linear regression equations. This data is not normalized by the off rate. **E.** Directly measured k_off_ of NINL from dynein at varied NINL concentrations in the presence (dark green) or absence (light green) of dynactin. The linear regression equation in the absence of dynactin is shown. **F.** Directly measured k_off_ of dynactin from dynein at varied dynactin concentrations in the presence (yellow) or absence (orange) of NINL. The linear regression equation in the absence of NINL is shown. **G.** Dwell times of NINL on dynein when NINL is labelled with TMR and JF646.Individual fluors represents the dwell times of these populations separately, and combined signal represents the dwell time if the signal from the two populations is combined into one channel. **H.** Intensity trace of NINL binding to dynein showing switching between the two fluorophores. **I.** Fraction of dynein bound to NINL at varied NINL concentrations in the presence of dynactin. Curves are fit to a single exponential. n ranges from 3-10. **J.** Fraction of dynein bound to dynactin at varied dynactin concentrations in the absence of NINL. Curves are fit to a single exponential. n ranges from 3-10.

**Supplement to Figure 3:**
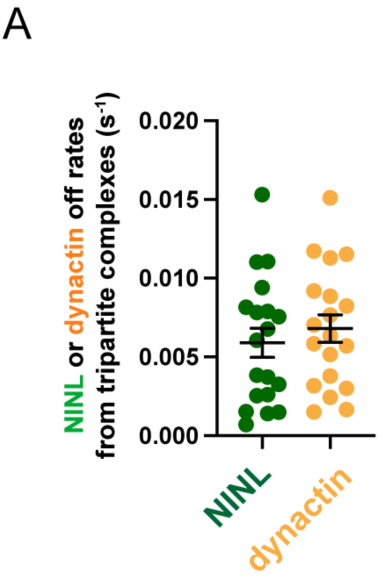
**A.** Directly measured k_off_ of NINL (green) or dynactin (yellow) from dynein-dynactin-NINL complexes.

**Supplement to Figure 4:**
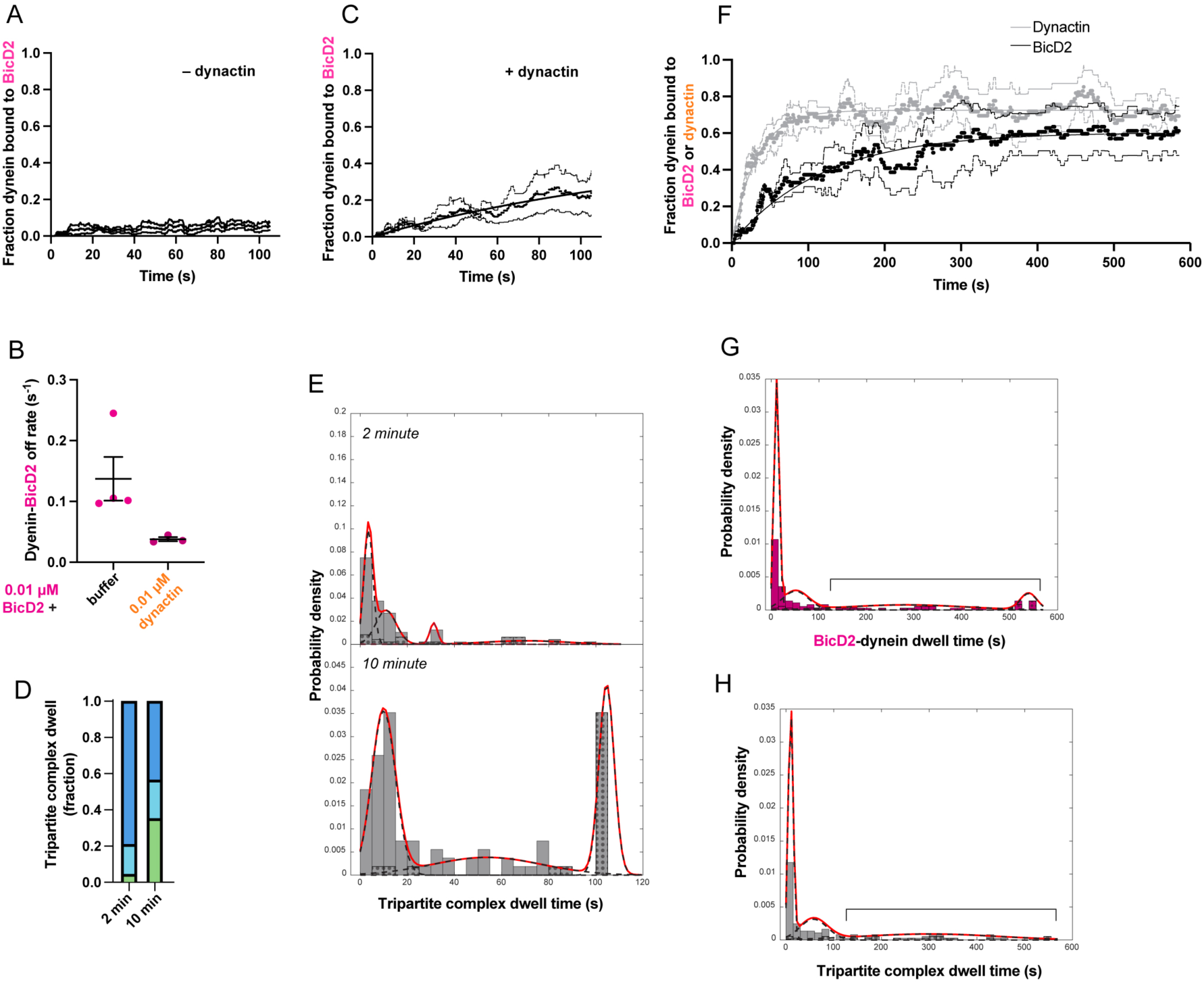
**A.** Fraction of dynein bound to BicD2 in the absence of dynactin. n = 4. **B.** Directly measured k_off_ of BicD2 from dynein in the presence and absence of dynactin. **C.** Fraction of dynein bound to BicD2 in the presence of dynactin over 2 min. Curve is fit to a single exponential. n = 3. **D.** Fraction of events from (E) that fall into short (<20s), medium (20-80s) or long (>80s) timescales. **E.** Histograms of the dwell time of dynein-dynactin-BicD2 complexes for imaging at 2 or 10 minutes. For the 10-minute data any dwell times greater than 110 seconds were randomly given a value between 100-110 and marked as censored for this histogram to better compare to the 2-minute data. Overall probability density function is shown as a solid red line, with individual component Gaussians as dashed black lines. Black dots show the percentage of the data in each bin that was truncated by the end of the acquisition. **F.** Fraction of dynein bound to BicD2 in the presence of dynactin over 10 min. Curve is fit to a single exponential. n = 3. **G.** Data from the bottom panel of (4A) without censoring events longer than 110s. Data that is subject to this censoring is indicated with the bracket. **H.** Data from the bottom panel of (S4E) without censoring events longer than 110s. Data that is subject to this censoring is indicated with the bracket.

